# Translational control of cardiac fibrosis

**DOI:** 10.1101/451666

**Authors:** Sonia Chothani, Sebastian Schafer, Eleonora Adami, Sivakumar Viswanathan, Anissa A Widjaja, Sarah R Langley, Jessie Tan, Chee Jian Pua, Giuseppe D’Agostino, Sebastiaan van Heesch, Franziska Witte, Leanne E Felkin, Eleni G. Christodoulou, Jinrui Dong, Susanne Blachut, Giannino Patone, Paul JR Barton, Norbert Hubner, Stuart A Cook, Owen JL Rackham

## Abstract

**Abstract:** *Background:* Fibrosis is a common pathology in many cardiac disorders and is driven by the activation of resident fibroblasts. The global post-transcriptional mechanisms underlying fibroblast-to-myofibroblast conversion in the heart have not been explored.

*Methods:* Genome-wide changes of RNA transcription and translation during human cardiac fibroblast activation were monitored with RNA sequencing and ribosome profiling. We then used miRNA-and RNA-binding protein-based analyses to identify translational regulators of fibrogenic genes. To reveal post-transcriptional mechanisms in the human fibrotic heart, we then integrated our findings with cardiac ribosome occupancy levels of 30 dilated cardiomyopathy patients.

*Results:* We generated nucleotide-resolution translatome data during the TGFβ1-driven cellular transition of human cardiac fibroblasts to myofibroblasts. This identified dynamic changes of RNA transcription and translation at several time points during the fibrotic response, revealing transient and early-responder genes. Remarkably, about one-third of all changes in gene expression in activated fibroblasts are subject to translational regulation and dynamic variation in ribosome occupancy affects protein abundance independent of RNA levels. Targets of RNA-binding proteins were strongly enriched in post-transcriptionally regulated genes, suggesting genes such as *MBNL2* can act as translational activators or repressors. Ribosome occupancy in the hearts of patients with dilated cardiomyopathy suggested an extensive post-transcriptional regulatory network underlying cardiac fibrosis. Key network hubs include RNA-binding proteins such as *PUM2* and *QKI* that work in concert to regulate the translation of target transcripts in human diseased hearts.

*Conclusions:* We reveal widespread translational effects of TGFβ1 and define novel post-transcriptional events that control the fibroblast-to-myofibroblast transition. Regulatory networks that affect ribosome occupancy in fibroblasts are paralleled in human heart disease. Our findings show the central importance of translational control in fibrosis and highlight novel pathogenic mechanisms in heart failure.

## Introduction

Cardiac remodelling and heart failure syndromes are frequently associated with fibrosis, which is a common late stage pathology in many human diseases ^1^. Cardiac fibrosis is seen in numerous cardiac conditions, including atrial fibrillation^2^, hypertrophic cardiomyopathy (HCM) ^3^, dilated cardiomyopathy (DCM)^4^ and heart failure with preserved ejection fraction^5^. Fibrosis of the heart is driven primarily by the activation of resident fibroblasts ^6, 7^. A better understanding of the molecular mechanisms underlying fibroblast activation is of great importance for the development of novel anti-fibrotic therapies ^8^.

While there are various cues known to initiate the cellular conversion of fibroblasts, Transforming growth factor β1 (TGFβ1) is considered the master regulator ^9^. However, anti-fibrotic therapeutic approaches based on TGFβ1 inhibition have side effects due to the pleiotropic roles of this cytokine, especially in cancer and inflammation^10,11^. Thus, unravelling the fibroblast-specific footprint of TGFβ1 signalling is an important step towards the identification of novel downstream drivers of cardiac disease. Whilst RNA expression changes via TGFβ1-induced SMAD signalling have been studied previously^12^, the independent impact of TGFβ1 on RNA translation remains unknown.

To address this gap in knowledge, we profiled genome-wide RNA transcription and translation levels^13^ in human primary cardiac fibroblasts at several time points after TGFβ1 stimulation. A tailored computational analysis^14^ identified post-transcriptional regulatory patterns underlying fibroblasts activation. To corroborate our findings in an independent and disease-relevant context, we performed ribosome profiling (Ribo-seq) of cardiac samples from patients with DCM. This integrative approach provided a detailed perspective on post-transcriptional regulatory hubs in human heart disease.

## Results

### Translational profiling during the activation of human cardiac fibroblasts

During the fibrotic response, resident fibroblasts become pro-fibrotic myofibroblasts that express α-smooth muscle actin (ACTA2) and secrete extracellular matrix (ECM) proteins such as collagen I and periostin (POSTN)^15^. To understand better this transition in the human heart, we isolated primary cardiac fibroblasts from atrial biopsies of four individuals undergoing coronary artery bypass grafting (**Supplementary table 1**). TGFβ1 stimulation (5ng/ml) resulted in a significant increase of ACTA2^+ve^ cells and an upregulation of ECM-related proteins indicating activation of fibroblasts within 24h (**Figure 1a-g**). This cellular transformation was accompanied by rapid phosphorylation of SMAD as well as ERK, which is a key factor in non-canonical signalling pathways and known to regulate post-transcriptional processes (**Figure 1h**)^12^.

**Figure 1:**
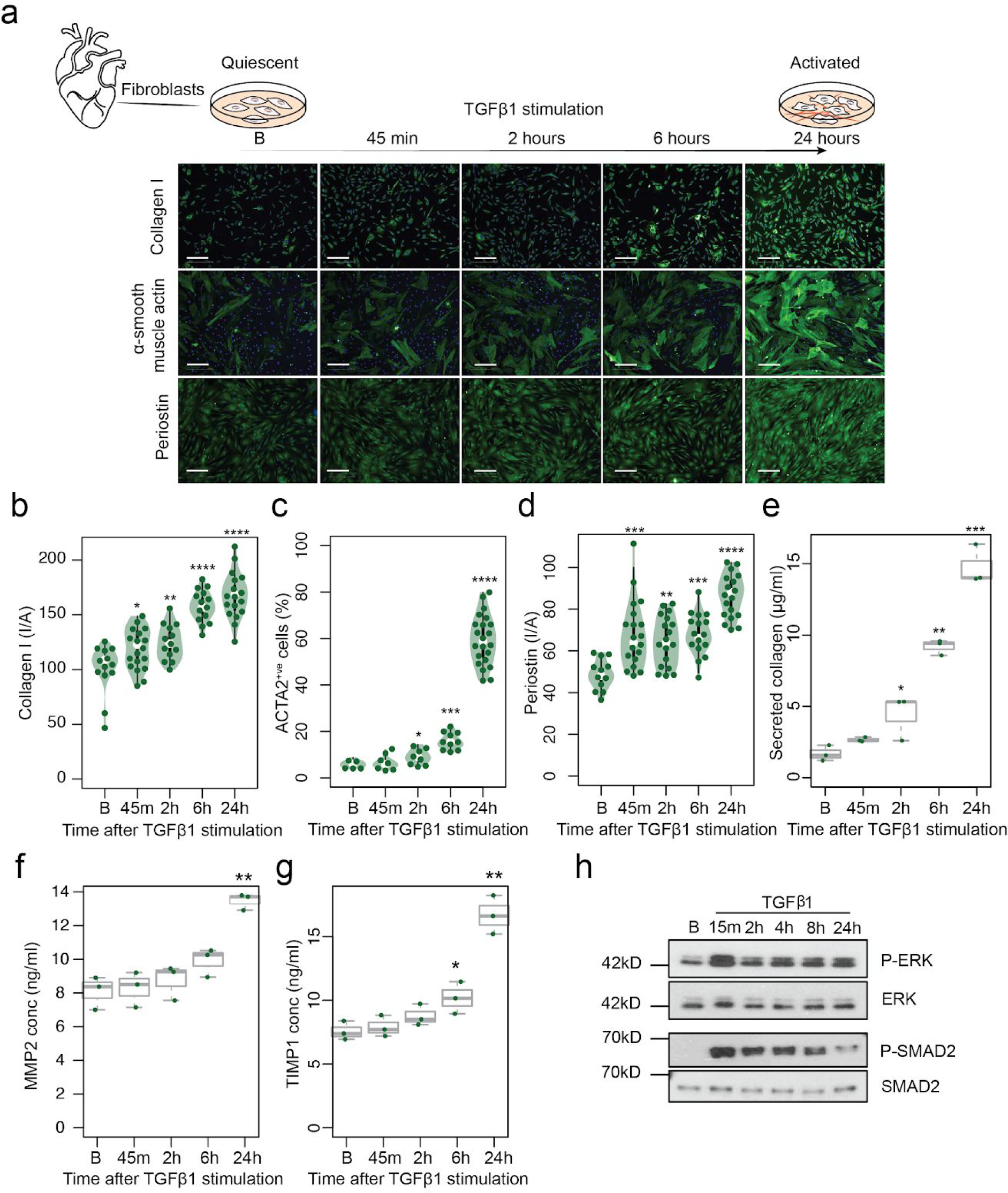
Time-resolved stimulation of the fibrotic response. **a.** Primary human cardiac fibroblasts were isolated from the atrial biopsies of four individuals and stimulated with TGFβ1 (5ng/ml). Microscopic images show fibroblasts at five timepoints (B: Baseline, 45m, 2h, 6h, 24h after TGFβ1 stimulation) with immunostaining for Collagen I, α-smooth muscle actin (ACTA2) and Periostin (POSTN). Scale bar equals 200µm. Fluorescence was quantified on the Operetta high-content imaging platform after immunostaining for Collagen I (**b**), ACTA2 (**c**) and POSTN (**d**) (28 measurements across four wells) and normalized for cell count (**c**) or cell area, I/A: Intensity/Area (**b**, **d**). Total secreted collagen (**e**), concentration of MMP-2 (**f**) and TIMP-1 (**g**) in the supernatant of TGFB1 stimulated cardiac fibroblasts (n=3, biologically independent samples) was quantified by Sirius red collagen assay (**e**) and by ELISA (**f, g**) respectively. P-values were determined by one-way ANOVA and corrected for comparisons to the same sample (Baseline) using Dunnett’s test. * p-value<5×10^−2^, ** p-value<10^−4^, *** p-value<10^−8^, **** p-value<2×10^−16^. **h.** Western blotting showed rapid activation of SMAD2 and ERK signalling molecules. B: baseline (0 minutes).

To capture a time-resolved snapshot of the molecular changes that underlie the transition of fibroblasts into myofibroblasts, we performed RNA∓seq and Ribo-seq for baseline, 45m, 2h, 6h and 24h after TGFβ1 stimulation. Ribo-seq entails deep sequencing of ribosomal footprints, which are short RNA fragments protected from nuclease treatment by ribosomes (RPF; ribosome protected fragments). RPFs therefore quantify both mRNA abundance and ribosome occupancy of protein-coding genes and as such are a superior proxy for protein levels compared to RNA-seq^16^. On average we generated ~51M (RNA-seq) and ~12M (Ribo-seq) uniquely mapped reads per sample. Ribo-seq reads mapped predominantly to the coding sequence and had an average read length of 29bp (**Supplementary Figure 1a-d**), both of which are characteristic of high-quality RPF data^13^. We then inferred the exact position of the Peptidyl-site (P-site), the site in the ribosome where transfer RNAs recognize their complementary codon, based on RPFs (see methods). On average, 89% of all P-sites mapped to the coding frame in known genes, indicating that captured ribosomes translate the known reading frame of transcripts (**Figure 2a, b**). Plotting of P-site density around the 3’ location of expressed CDS revealed that ribosomes recognised the stop codon and disengaged from RNA transcripts (**Figure 2c**). High triplet periodicity and a strong dissociation signal reveal the stepwise movement of the ribosome along the coding regions of the transcripts and indicate the positions of actively translating ribosomes being captured at single nucleotide resolution.

**Figure 2:**
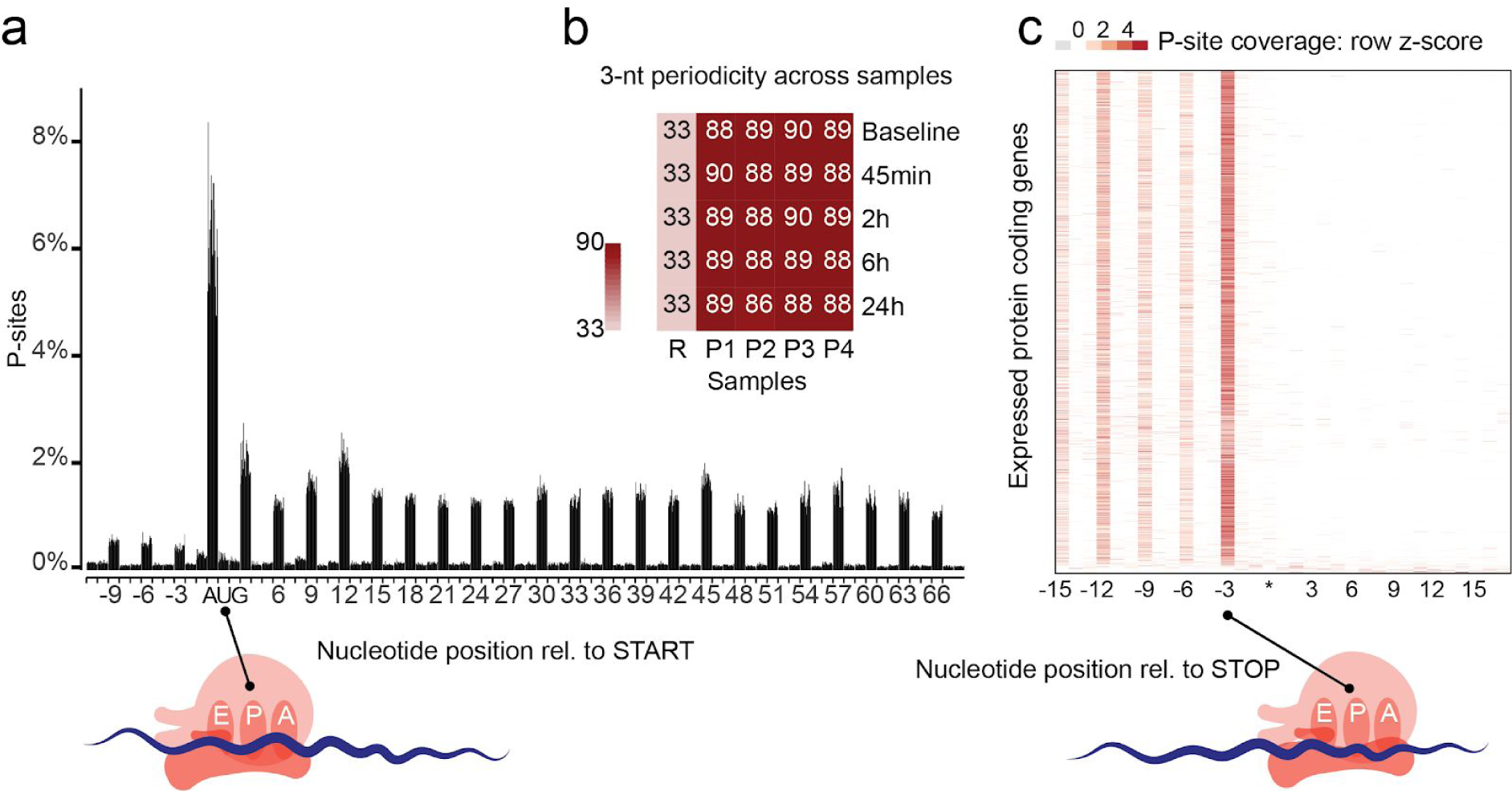
Ribosome profiling of TGFβ1 stimulated primary human cardiac fibroblasts. **a**. Sample level periodicity: Distribution of inferred P-site (Peptidyl-site) locations (+12 offset) for each sample (4 patients over 5 time points) at annotated translation start sites reveals ribosomes located on the canonical start codon (AUG) and majority of the P-sites downstream of the start codon located in-frame. **b.** The 3nt-periodicity for all 20 samples (P1-P4 patients, 5 time points) is >86% indicating the majority of reads represent actively translating ribosomes. R represents random 3nt-periodicity of 33%. **c.** Gene-level periodicity: P-site location across all annotated expressed (Transcript Per Million mapped reads, TPM>1) genes (combined data from 4 patients over 5 time points), shows efficient ribosome drop-off at the canonical stop codon (UGA/UAG/UAA, represented by *).

### Dynamic translational regulation during the fibrotic response

Differentially transcribed genes (DTGs) can be detected on a genome-wide scale with RNA sequencing alone. Conversely, genes that are translationally regulated between conditions will display a significant change in translational efficiency (TE), i.e. the gene-wise ratio between ribosome occupancy and transcript abundance, requiring both RNA-and Ribo-seq for detection. We recently developed an analytical approach (which we refer to as ΔTE) that integrates RNA-seq and Ribo-seq data to reveal differential translational-efficiency genes (DTEGs)^14^. An output of this approach is a ΔTE value (and associated adjusted P-value) for each gene describing the log fold-change of TE at each time point. This analysis allowed reliable detection of DTEGs in our data despite patient-related batch effects (**Supplementary Figure 2**). Using the ΔTE approach we identified 1,691 DTEGs during the fibrotic response. For instance, ribosome occupancy of both *FTL (Ferritin Light Chain*, ΔTE=3.24; P_adj._=2×10^−2^) and *FTH1* (Ferritin Heavy Chain 1, ΔTE=3.12; P_adj._=1×10^−2^) increased significantly upon TGFβ1 stimulation, despite underlying transcript levels remaining the same. Translating ribosomes located on *ITGA3* (*Integrin Subunit Alpha 3,* ΔTE=−1.9; P_adj._=2×10^-3^) transcripts decreased despite constant levels of RNA. These dynamic and often transient post-transcriptional changes in gene expression were sufficient to affect protein expression (**Figure 3a**, **Supplementary Figure 3a-d**). Globally, TGFβ1 signalling had an immediate effect on the ribosome occupancy of 67 genes after 45 minutes (**Figure 3b**). The most enriched gene ontology (GO) term in these early responding genes was “transcription regulator activity” (P_adj._= ×10^−3^), suggesting that the following transcriptional response may be modulated by these DTEGs. The impact on translation then gradually decreased at 2h and 6h but was very pronounced again at 24h (**Supplementary File 1**, **Supplementary Figure 3g-j**).

**Figure 3:**
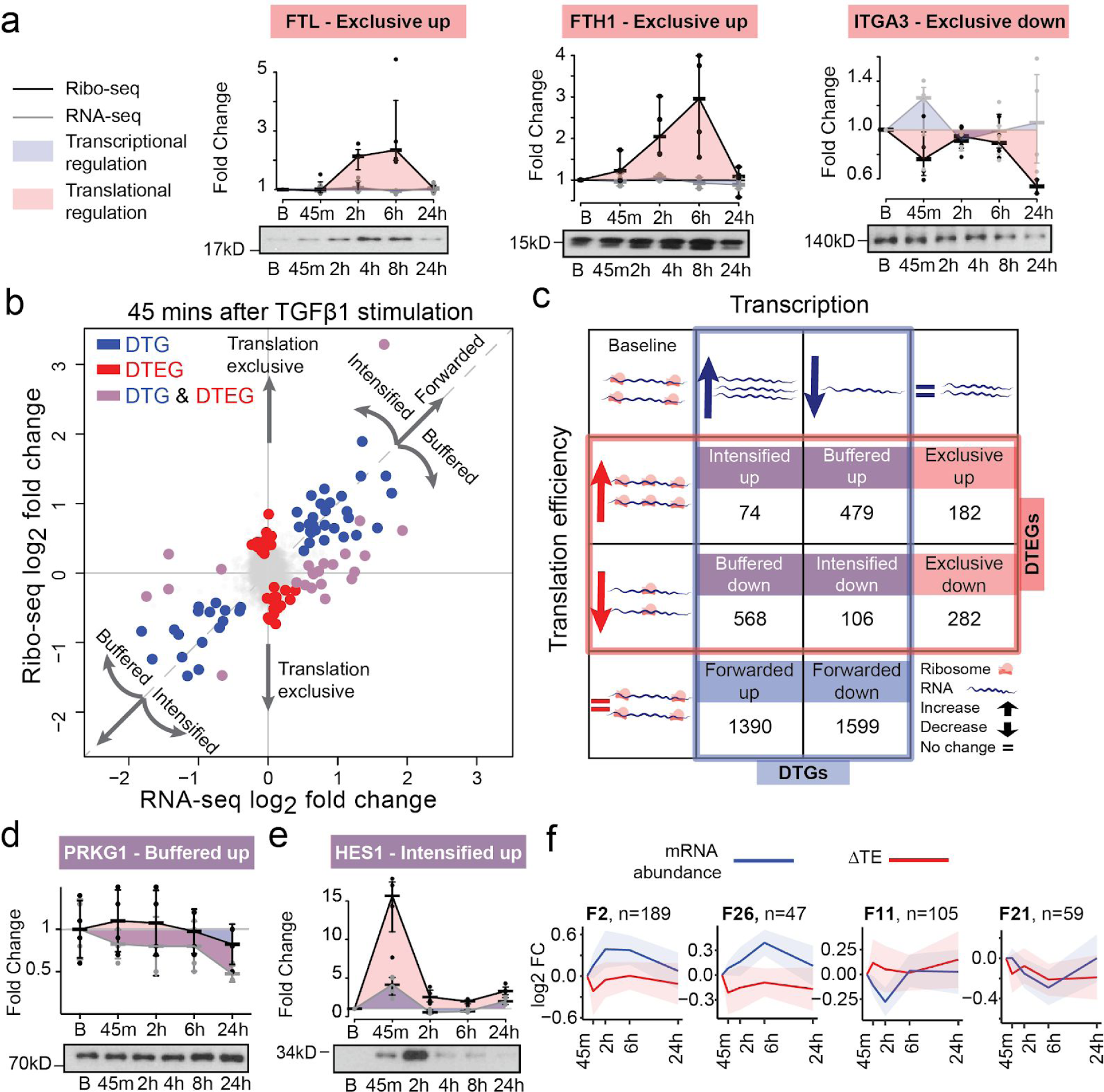
Genome-wide temporal transcriptional and translational landscape in cardiac fibrosis. **a.** Western blots showing ribosome occupancy determining changes in protein levels independent of mRNA changes for translationally exclusive genes, *FTL*,*FTH1*,*ITGA3*. B: Basal. **b.** Log-fold changes in the mRNA and ribosome occupancy at 45 minutes after TGFβ1 stimulation. DTG: Differentially transcribed genes, DTEG: Differential translational-efficiency genes. **c.** The interplay between DTGs and DTEGs showing several categories of gene expression regulation. Forwarded genes, where the occupancy changes are explained by the mRNA changes; Exclusive, where changes occur exclusively in TE without underlying mRNA changes; Buffered and Intensified, where both the TE and the mRNA are changing.**d-e.** Western blotting of genes detected as **d.** buffered, PRKG1 and **e.** intensified, HES1. **f.** Forwarded gene clusters (F2, F26, F11, F21) with transient changes in expression following TGFβ1 stimulation. n: Number of genes in the cluster, FC: fold change.

We also detected a total of 4,216 DTGs, which gradually increased across time points. Several genes were detected as both DTG and DTEG; This occurs when TGFβ1 affects both RNA levels and TE of the same gene. In order to describe the relationship of this overlap in transcriptional and translational regulation, we categorise each of the DTGs and DTEGs into one of eight regulatory groups (**Figure 3c**, **Supplementary File 2**). For more than 29% of DTGs, differences in transcription were not forwarded to the translational level but were translationally buffered or intensified. Of these, translational buffering was most prominent, i.e. changes in transcript expression detected with RNA-seq were less pronounced in the Ribo-seq data. This effect can be due to either a less efficient translation of genes whose RNA levels are increasing (568 genes, buffered down) or vice-versa, more efficient translation of genes whose RNA levels are decreasing (479 genes, buffered up). In particular, out of these 1,047 buffered genes, 419 (231 down, 188 up) had transcriptional regulation that was completely counter-acted by translational regulation, resulting in a similar density of translating ribosomes despite underlying TGFβ1-driven transcriptional changes. For example, RNA-seq suggests a downregulation of the protein kinase PRKG1 upon TGFβ1 stimulation. However, this effect is not forwarded to the translational level and thus protein levels do not decrease (**Figure 3d**, **Supplementary Figure 3f**).

For a defined subset of transcripts, RNA expression differences were intensified at the level of translation during the fibrotic response. These genes (n=180) responded even more strongly to TGFβ1 treatment than would be expected from RNA-seq based analyses alone. For instance, the concerted upregulation of the transcription factor *HES1* on both the transcriptional and translational level resulted in a very strong increase in RPFs, which resulted in a profound increase in HES1 protein (**Figure 3e**, **Supplementary Figure 3e**). Interestingly, intensified genes were over-represented for functions such as “SMAD-protein signal transduction” (P_adj._=6.5×10^-3^) and “Regulation of ERK1 and ERK2 cascade” (P_adj._=1.4×10^-2^) (see **Supplementary file 3**).

Overall these results demonstrate that more than one-third of all gene expression changes during fibroblast activation involve translational regulation and that these changes can affect protein levels.

Transcriptional and translational regulation downstream of TGFβ1 appears to be closely interlinked and tightly regulated over time. To further stratify these effects over time, we performed unsupervised clustering of the temporal profiles of Forwarded, Buffered, Exclusive and Intensified genes. This revealed 64 distinct regulatory patterns during the fibrotic response (**Supplementary Figure 4**). The clustering highlights the fact that there are a substantial number of transiently regulated genes during the fibrotic response (**Figure 3f**). Transient differences in expression would not be apparent when quiescent fibroblasts are compared to myofibroblasts at 24h, but may be crucial for the cellular transition and therefore important in disease. Transient clusters were predominantly enriched for processes involved in the regulation of gene expression(regulation of RNA metabolic process, 2.5×10^−7^; regulation of transcription, DNA-templated, 1.7×10^−5^), further substantiating their role in fibroblast transformation (**Supplementary File 4**).

### Post-transcriptional regulators of TGFβ1 signalling

Having identified different regulatory groups and temporal profiles we sought to identify possible regulators. It is known that both RNA binding proteins (RBPs) and micro-RNAs (miRNAs) bind to target transcripts and affect protein production^16,17^, a number of which been previously linked to heart disease^18,19^. To identify key post-transcriptional regulators during the fibrotic response, we integrated expression data of possible regulators (miRNAs/RBPs) with transcriptome-wide target binding data. We performed global miRNA sequencing at Baseline, 45m, 2h, 6h and 24h after TGFβ1 stimulation in human primary cardiac fibroblasts (n=4), revealing 15 differentially expressed miRNAs (**Supplementary Figure 5**, **Supplementary Table 2**). For each of these miRNAs, a permutation test was used to detect a significant over-representation of their targets (‘Very High’ confidence, mirDip^20^) in each of the eight regulatory groups defined above (**Figure 4a**). Targets of downregulated miRNAs miR-3182 (P_adj._=0.0364), miR-335-3p (P_adj._=0.0024) and miR-101-3p (P_adj._=4.9×10^-05^) were significantly enriched mainly in the ‘buffered up’ group (**Supplementary File 5**). However, the overall role of miRNA regulation appears to be limited in the TGFβ1-driven fibroblast-to-myofibroblast transition.

**Figure 4:**
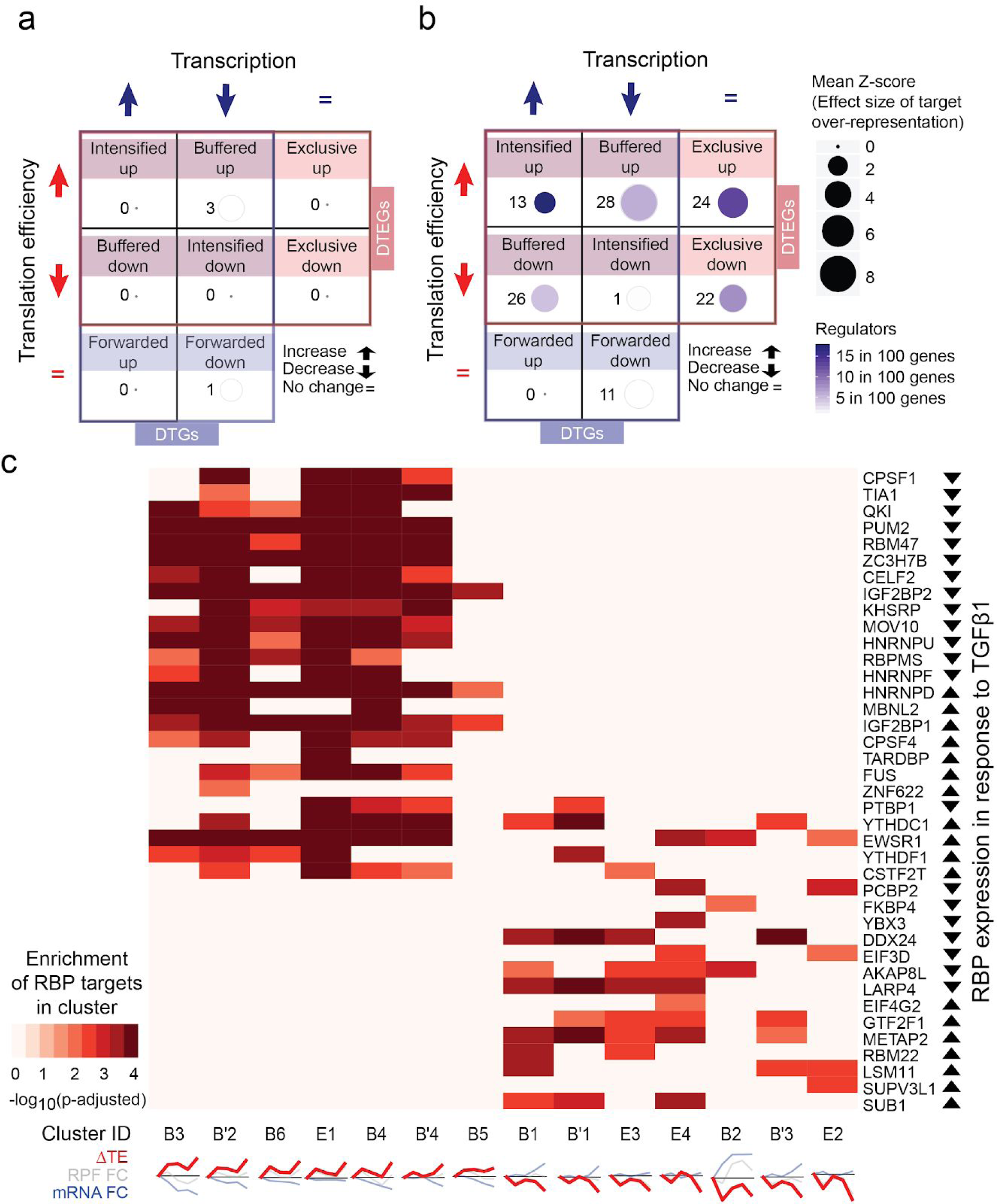
Post-transcriptional regulators in fibroblast activation. **a.** miRNA and **b.** RBP target over-representation test in the regulatory groups within DTGs and DTEGs. Z-score is the effect size of over-representation. Regulators per member represent the number of miRNAs or RBPs over-represented per member of the group. **c.** RBP over-representation test (FDR<1%) for regulatory patterns separate translationally activated and repressed clusters. RBP expression in response to TGFβ1 is determined using significant RPF changes. RBPs that are not over-represented for their targets in any cluster are not shown. Cluster IDs are denoted by their regulatory groups and cluster number. B: Buffered, B’: Completely buffered (special case) and E: Exclusive. Clusters with less than 50 genes, or with no RBP over-representation are not shown.

Using the same methodology, we tested for an overrepresentation of RBP targets in differentially expressed genes during the activation of fibroblasts. To construct reliable RBP-target networks, we utilized experimental evidence for RBP-RNA binding derived from eCLIP (n=117), PAR-CLIP^21^ (n=58), HITS-CLIP (n=23) and iCLIP (n=22) data provided by ENCODE^22^ and POSTAR^23^. RBP targets were predominantly enriched in DTEGs but not in DTGs, which suggests that RBPs shape the fibrotic response mainly through translational regulation and rarely influence transcript levels (**Figure 4b**). Most targets of individual RBPs were overrepresented in clusters with similar translational regulation profiles (**Figure 4c**). Distinct enrichments in unidirectional ΔTE patterns suggest RBPs tend to act either as translational repressors or activators during the fibrotic response.

### Translational regulation by RBPs in fibrotic DCM heart

We reported previously an in-depth view of the cardiac transcriptome in health and DCM^24^. Fibrotic markers^6,25^ were upregulated in the hearts of end-stage DCM patients (**Figure 5a**). In total, 45 of the 47 RBPs identified previously were also detected in DCM hearts (Transcripts per million mapped reads, TPM>5) and 22 were differentially expressed between healthy and diseased individuals (FC≧|1.2|, P_adj._≦0.05) (**Figure 5b**). To quantify translation levels in the fibrotic human heart, we performed ribosome profiling of left ventricular tissue collected from a subset of DCM patients. In total, we were able to generate high depth and high-quality Ribo-seq data of 30 individuals (**Supplementary Figure 6**, see methods for details of data generation).

**Figure 5:**
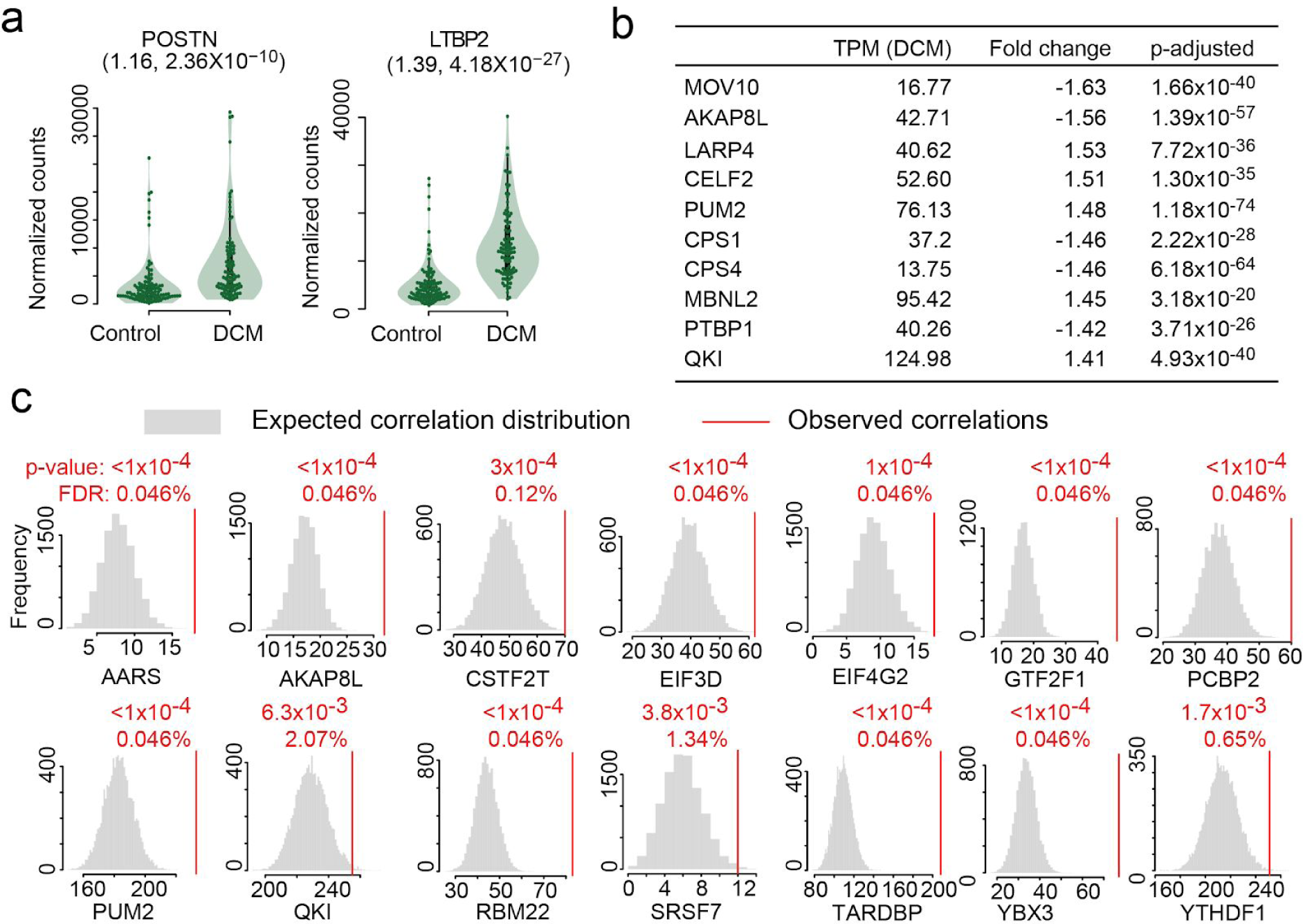
Post-transcriptional regulators in fibrosis and dilated cardiomyopathy. **a.** Periostin (POSTN) and Latent TGFB binding protein 2 (LTBP2) are upregulated in DCM patients. (Fold change, FDR-adjusted P value) **b.**Cardiac expression (Transcripts per million mapped reads, TPM) and differential expression (fold change, p-adjusted: p-value corrected by Benjamini-Hochberg) of RNA binding proteins in DCM patients compared to non-diseased donors. **c.** RBPs with significantly more correlated targets than expected by chance (RBP RPF vs target TE, p-values are nominal, FDR are corrected for False Discovery Rate by the Benjamini-Hochberg method), indicating translational control also in DCM patients.

**Figure 6:**
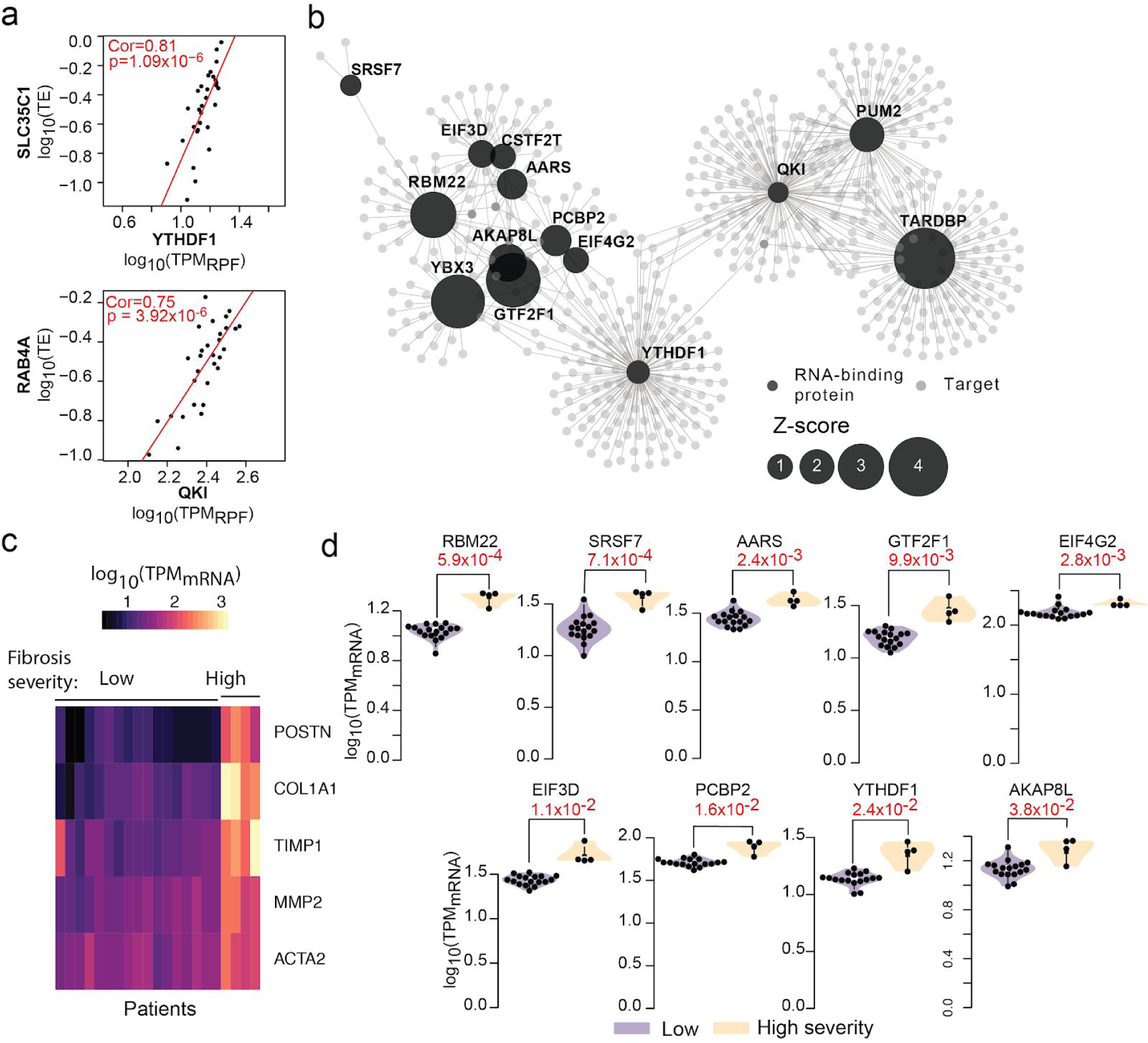
Post-transcriptional regulatory network in dilated cardiomyopathy. **a.** Exemplars of RBP-target pairs correlated in the DCM heart. Cor: Spearman ranked correlation value, p: p-value for correlation test. **b.** RBP-target network in disease based on permutation tests (|ρ| ≥ 0.45 for visualisation). **c.** Patient stratification based on severity of fibrosis assessed using marker gene expression (only low and high severity groups shown, full clustering in supplementary figure 8b) **d.** RBP expression differences between patients with low and high severity of fibrosis. In red, p value for two-tailed t-test.

### Post-transcriptional regulatory networks in DCM patients

Genetic variation and differences in disease severity contribute to varied gene transcription and translation levels in the diseased heart. If the RBPs are truly regulators of ribosome occupancy in fibrosis, then RBP expression levels should correlate with the translational efficiency of the target transcripts. We observed a total of 3,771 RBP:target pairs correlating significantly in our human heart datasets. Fourteen translational regulators (targeting 926 transcripts) remained significantly correlated with their targets based on permutation analyses (**Figures 5c, 6a,b**, **Supplementary File 6**), identifying them as regulatory hubs controlling ribosome occupancy during cardiac fibroblast activation and in the hearts of DCM patients. This substantiates further the influence of RBPs on ribosome occupancy in an independent dataset and provides evidence for post-transcriptional regulatory networks in human heart.

Modules of this post-transcriptional regulatory network are enriched for distinct biological functions related to actin remodelling and other processes important in fibrosis (**Supplementary File 7**). One module in the heart is controlled by *PUM2*, which is known to drive glial scar formation, a fibrotic and TGFβ1-dependent process in the brain^26^. Others are regulated by QKI, important in heart development^27^, or *PCBP2* which is known to inhibit cardiac hypertrophy driven by the fibrogenic stimulus Angiotensin 2^28^. Interestingly, we detected a large overlap between targets of regulators, suggesting that the concerted action of several RBPs determine the translational efficiency of bound transcripts. Both *QKI* and *PUM2* appear to cooperate, with more than 21% of their targets (103 genes) overlapping between the RBPs, both of which appear to be acting as translational repressors.

To test whether the differentially regulated RBPs we discovered in the DCM samples contribute to cardiac fibrosis, we determined whether the expression of these RBPs correlated with the same fibrosis markers used in figure 1. All fourteen identified regulatory hubs were correlated significantly with at least one marker gene such as *ACTA2*, *COL1A1* or *POSTN* (**Supplementary Fig. 7a**), which supports their involvement in fibrosis in the DCM heart. Unsupervised clustering revealed different degrees of fibrogenic marker gene expression (low, moderate, high) in the DCM hearts (**Figure 6c**, **Supplementary Fig. 7b**). Nine regulators of the network were significantly elevated in patients with high severity (vs low severity) cardiac fibrosis molecular signature (two-tailed t-test significance p-value<0.05, **Figure 6d**).

## Discussion

Here we show that TGFβ1 stimulation of human atrial fibroblasts causes rapid changes in translation of a distinct set of key fibrosis transcripts, some of which occur within minutes after stimulation. While the transcriptional effects of TGFβ1 are well studied^12^, we show for the first time a genome-wide snapshot of the post-transcriptional effect of the pathway: over one-third of all changing genes are regulated at the translational level. Buffering or intensifying RNA differences and exclusive translational regulation robustly modify protein abundance during the cellular transformation to myofibroblasts. These results shed some new light on the extent of post-transcriptional regulation in response to extracellular signalling in human disease.

We found that targets of specific miRNAs or RBPs are significantly enriched in translationally but not transcriptionally regulated genes, revealing their post-transcriptional regulatory footprint during cardiac fibroblast activation. miR-101 was previously reported to inhibit cardiac fibrosis via the TGFβ1 pathway^29^, block fibroblast activation^30^ and is reduced in the lung and serum of IPF patients^31^. We show that TGFβ1 stimulation results in a loss of anti-fibrotic miRNA-101 in atrial fibroblasts, which is significantly associated with an increase in ribosome occupancy on a select number of target transcripts (**Supplementary File 5**). These observations are consistent with the recognized role of miRNAs in the repression of translation^32^.

Integration of more than 150 protein-RNA binding data sets with our ΔTE analysis revealed an unprecedented view of the RBP-driven landscape of translational regulation. *CELF2^33^*, *PUM2^26^* and *KHSRP^34^* are known to repress whereas *MBNL2^35^* and *LARP4^36^* to activate the translation of single target genes. Our results expose their interrelated larger target network in fibrotic disease and confirm they predominantly function as either translational repressors or activators, respectively. Multi–functional RBPs can regulate many post-transcriptional processes such as mRNA splicing, localization, translation and turnover in parallel^37^. Quaking (QKI) and MBNL2 are mostly associated with alternative splicing^38–40^, but our findings suggest that both also regulate the translational efficiency of hundreds of target transcripts. Some isoforms of QKI shuttle between the nucleus and the cytoplasm during the fibrotic response. This is likely related to their role in the subcellular localization of target transcripts^40^. Some isoforms of QKI shuttle between the nucleus and the cytoplasm^41^ and have been shown to regulate the translation of luciferase reporter RNAs^42^.

We documented highly significant correlation of fourteen RBPs with the translational efficiency of more than 900 transcripts in the DCM heart and identified key regulatory hubs of the diseased cardiac translatome. Regulatory footprints of the network often overlap, suggesting that RBPs act in concert to control the final protein levels of shared targets. This is especially evident with the repressors *PUM2* and *QKI* with more than 21% of their targets overlapping. Translational repression by *PUM2* in astrocytes is known to cause astrogliosis and the formation of glial scarring, which is a TGFβ1 dependent process^26^ and we reveal here the target gene network for *PUM2*–regulated translation in cardiac scarring.

More than 1,500 RBPs are encoded in the human genome^43^, but their role in the translation of target mRNAs remains largely unexplored. From our studies it is apparent that there is a major regulation of cardiac fibroblast gene expression at the level of translation and that this level of control, which is equally important as transcriptional regulation, has not been appreciated in this context. Translational control may be particularly pertinent in fibroblasts that produce huge amounts of ECM protein with limited changes in the expression of the correspondingly ECM transcript^44^. Whether such processes are important in cardiac myocytes or other cells in the heart is still to be determined, but their role in the regulation of disease is clear, opening the possibility of alternative targets for treatment.

## Methods

### Human primary fibroblast culture

Human atrial fibroblasts were prepared and cultured as described previously^12^. All the experiments were carried out at low cell passage (<P4). In all experiments, cells were starved in serum-free DMEM for 16 hours prior to TGFβ1 stimulation. Stimulated fibroblasts were compared to unstimulated fibroblasts that have been grown for the same duration under the same conditions (serum-free DMEM), but without the stimuli.

### Operetta high-content imaging

Operetta phenotyping assay was performed as described previously^12^. Briefly, atrial fibroblasts were seeded in 96-well black CellCarrier plates (PerkinElmer) at a density of 10^4^ cells/well. Following TGFβ1 stimulation, cells were fixed in 4% paraformaldehyde (28908, Life Technologies), permeabilized with 0.1%Triton X-100 (Sigma) and non-specific sites were blocked with 0.5% BSA and 0.1% Tween −20 in PBS. Cells were incubated overnight (4°C) with the following primary antibodies (1:500): alpha-smooth muscle actin (ACTA2, ab7817, abcam), Collagen I (ab34710, abcam) and periostin (POSTN, ab14041, abcam), followed by incubation with the appropriate AlexaFluor 488 secondary antibodies (1:1000), and counter-staining with rhodamine-phalloidin (1:1000, R415, Life Technologies) and DAPI (1µg/ml, D1306, Life Technologies). Each condition was imaged from at least two wells and a minimum of 7 fields per well using Operetta high-content imaging system 1483 (PerkinElmer). The quantification of ACTA2 positive cells were measured using Harmony v3.5.2 (PerkinElmer). The measurement of fluorescence intensity per area of Collagen I and POSTN fluorescence intensity per area (normalized to the number of cells) were performed with Columbus 2.7.1 (PerkinElmer).

### Enzyme-linked immunosorbent assay (ELISA)

The amount of MMP-2 and TIMP-1 in equal volume of cell culture media was quantified using Total MMP-2 (MMP200, R&D Systems) and TIMP-1 Quantikine ELISA kit (DTM100, R&D Systems) as per the manufacturer’s instructions.

### Colorimetric Assay

Quantification of total secreted collagen in the cell culture supernatant was performed using Sirius red collagen detection kit (9062, Chondrex) according to the manufacturer’s protocol.

### Western blotting

Atrial fibroblasts were washed with ice-cold PBS and solubilized by gentle rocking in radioimmunoprecipitation assay (RIPA) buffer containing protease and phosphatase inhibitors (Roche). Protein concentrations were determined by Bradford assay (Bio-Rad). After centrifugation, equal amounts of protein lysates were separated by SDS-PAGE, transferred to PVDF membrane, and subjected to immunoblot analysis for: p-ERK1/2 (4370, CST), ERK1/2 (4695, CST), FTL (ab109373, Abcam), FTH1 (4393, CST), GAPDH (2118, CST), HES1 (11988, CST), Integrin α3 (ITGA3, sc-374242, SantaCruz), PKG-1 (3248, CST), p-SMAD2 (5339, CST), SMAD2 (3108, CST). Proteins were visualized using the ECL detection system (Pierce) with the appropriate secondary antibodies.

### Statistical analysis

Statistical analyses of high content imaging was performed using GraphPad Prism software (version 6.07). Outliers (ROUT 2%, Prism Software) were removed before analysis. Dunnett’s test was used to calculate multiple testing corrected p-values for comparison of several time points to baseline.

### Ribosome profiling and RNA sequencing

#### Primary cells

Primary human atrial fibroblasts outgrown from cardiac tissues biopsies of four patients undergoing coronary artery bypass grafting were expanded to reach 80% confluency in several 10cm dishes. Cells were stimulated with 5 ng/ml TGFβ1 for 45 min, 2h, 6h and 24h. Per condition, three 10cm dishes were employed in order to obtain enough material for Ribo-seq (two dishes) and RNA-seq (one dish).

#### Heart tissue

As part of a larger consortium effort (manuscript in preparation) to characterize the cardiac translatome, we generated ribosome profiling data of left ventricular tissue samples collected during left ventricular device implantation or cardiac transplantation from patients with end-stage DCM (n = 30). We specifically selected these 30 DCM patients as they were obtained from the same site of tissue collection (Cardiovascular Research Centre Biobank at Royal Brompton and Harefield NHS Trust.), in order to reduce technical variability and facilitate accurate patient stratification based on the degree of cardiac fibrosis.

Ribosome profiling was performed as previously described^16^. Briefly, snap-frozen cell pellets or 50-100 mg of tissue, previously powdered under liquid nitrogen, were lysed in 1ml cold lysis buffer (formulation as in TruSeq Ribosome Profile, Illumina) supplemented with 0.1mg/ml cycloheximide (CHX) to stabilize ribosomal subunits and prevent post-lysis translocation. Homogenized and cleared lysates were then footprinted with Truseq Nuclease (Illumina) according to manufacturer’s instructions. Ribosomes were purified using Illustra Sephacryl S400 columns (GE Healthcare) and the protected RNA fragments were extracted with standard phenol:chloroform:isoamylalcohol technique. Following ribosomal RNA removal (Mammalian RiboZero Magnetic Gold, Illumina), sequencing libraries were prepared out of the footprinted RNA. Ribo-seq libraries were pooled to perform multiplex sequencing on Illumina Hiseq machines.

The RNAseq data for the heart tissue was derived from Henig et al^24^. To prepare polyA+ RNA-seq libraries from primary cardiac fibroblasts, total RNA was extracted with Trizol from one 10cm dish per condition. Following cleanup with RNeasy kit (Qiagen), ~500ng of each sample were further processed with the Truseq Stranded mRNA kit (Illumina). Barcoded RNA-seq libraries were pooled and sequenced on the Illumina HiSeq platform.

#### Informed Consent

DCM tissue studies complied with UK Human Tissue Act guidelines and were carried out with approval from the Royal Brompton and Harefield local ethical review committee and the National Research Ethics Service Committee South Central, Hampshire B (reference 09/H0504/104)

## Data processing for RNA-sequencing and Ribosome profiling

Raw sequencing data were demultiplexed with bcl2fastq *V2.19.0.316* and the adaptors were trimmed using *Trimmomatic^45^ V0.36*, retaining reads longer than 20 nt post-clipping. RNA-seq reads were further clipped with FASTX Toolkit *V0.0.14* to 29nt, to allow 14 comparison directly with Ribo-seq reads. Reads were aligned using bowtie^46^ to known rRNA, tRNA and mt-rRNA sequences (RNACentral^47^, release 5.0), aligned reads were filtered out to obtain only RPFs. Alignment to the human genome (hg38) was done using STAR^48^. Gene expression was quantified on the CDS (coding sequence region) using uniquely mapped reads (Ensembl database release GRCh38 v86 combined with additional transcripts from RefSeq GRCh38, latest version downloaded January 2018) with feature counts^49^. Genes with mean transcripts per million mapped reads, TPM < 1 in either RNA-seq or Ribo-seq across all conditions were removed before downstream analysis. Ribotaper^50^ was used to obtain the in-frame reads around the start and stop codon. These p-sites were then visualized across samples and genes. Heatmap for the ribosome drop-off was generated using pheatmap *1.0.8* R package^51^. Data quality check was done using MultiQC^52^. Principal component analysis was carried out using prcomp function in R. Individual gene batch effect for ITGA7 was removed using limma package^53^ in R (see details in **Supplementary table 1**).

## Detection of Differential translational-efficiency genes (DTEGs) using DESeq2

The calculation of ΔTE for human fibroblast data was done using an interaction term while accommodating for patient effect in the statistical model within DESeq2^54^ (*~ Patient + Time + Sequencing + Time:Sequencing*). This allows for the identification of significant differences between time-points that are discordant between sequencing methodologies, i.e. for changes in ribosome occupancy that are not explained by changes in RNA abundance. The ΔTE fold change derived with this approach is comparable to traditional TE, but also accounts for variance and level of expression. In addition, the statistical model reveals if ΔTE is statistically significant. Since the RNA and RPF fold changes can be obtained by the same process using DESeq2, the fold changes are also directly comparable with ΔTE. In combination, these three fold changes can help predict the regulation status of the gene at transcriptional and translational level. Results were combined from Wald test for each time point and likelihood ratio test across all time points. Published tools for DTEG detection were run using default parameters (**Supplementary table 4**).

## Classification of DTGs and DTEGs into regulatory classes

A gene’s regulation class was inferred using significant p-adjusted value threshold of 0.05 for ΔTE, differential expression of RPF and RNA, as shown in **Supplementary table 3**. Forwarded genes, change at the RPF and RNA levels but do not have a significant change in TE. Exclusive genes had a significant change in RPFs and TE, but no change on the mRNA levels. Buffered and Intensified genes had all three levels significantly changing, but the relative direction of RNA and TE changes were used to determine the gene’s regulation status. Buffered genes were ones where the mRNA changes were counteracted by TE changes. Hence, genes with fold changes of RNA and TE in opposite directions were considered buffered. A special case of buffered genes, were significant at the RNA levels, with a counteracting significant change in TE, and no significant change in RPF indicating complete translational buffering. Intensified genes had TE changing in the same direction as RNA, hence reinforcing the transcriptional effect. The changes that were significant were used for hierarchical clustering within each category of DTEGs and DTGs. For instance forwarded genes were clustered using the RPF and mRNA fold changes. Hierarchical clustering was carried out using euclidean distances and ward.D method in hclust function^55^ in R stats package. Clusters were obtained using cutreeDynamic^56^ with default settings.

## miRNA sequencing

Purified RNA was quantified using Qubit RNA high sensitivity assay kit (Life Technologies) and RNA integrity number (RIN) was measured using the Bioanalyzer RNA 6000 Nano assay (Agilent Technologies). TruSeq Small RNA Library Preparation Kit (Illumina) was used to assess miRNA abundance following manufacturer’s protocol. Briefly, 3’ and 5’ ends of 50ng purified RNA with RIN value >6 was ligated with RNA adaptors before reversed transcribed into cDNA and amplified. The amplified cDNA constructs were resolved using denaturing PAGE purification in Novex TBE 6% gel (Thermo Fisher Scientific). Targeted cDNA constructs with size between 145-160bp containing ~22nt mature miRNA were recovered and concentrated using ethanol precipitation. The final libraries were quantified using KAPA library quantification kits (KAPA Biosystems) on StepOnePlus Real-Time PCR system (Applied Biosystems) according to manufacturer’s guide. The quality and average fragment size of the final libraries were determined using LabChip GX DNA High Sensitivity Reagent Kit (Perkin Elmer). Libraries with unique indexes were pooled and sequenced on a NextSeq 500 benchtop sequencer (Illumina) using NextSeq 500 High Output v2 kit and single-end 50bp sequencing chemistry. 4SeqGUI, docker4seq reproducible pipeline was used to process the miRNA-seq^57^. DESeq2 was used for differential expression analysis, using patient as a covariate. Results were combined from Wald test and likelihood ratio test.

## Over-representation analysis

R package topGO^58^ and KEGGrest^59^ were used to carry out (GO: BP, MF and KEGG pathways) over–representation tests for each gene cluster. Genes that are classified as either DTG or DTEG were used as background. miRNA-target enrichment was carried out for miRNA target data (for miRNAs with CPM > 1) downloaded from mirDIP using the ‘Very High’ confidence score class^20^. For each miRNA, actual number of targets in each gene class or cluster were determined using the database. Expected distribution for miRNA targets in each group was calculated by randomly selecting gene sets of the same size and quantifying the targets found within the group. This was repeated 100,000 times to obtain an empirical p-value. These p-values were further corrected for multiple testing using *Benjamini-Hochberg* method. The groups with BH corrected p-value < 0.05 for a miRNA were considered over–represented for its targets. Z-scores were calculated to evaluate the effect size of this over–representation. RBP’s target over-representation was also identified in regulatory groups using a similar permutation analysis, but using CLIP-seq data for target binding. Peak files from eCLIP experiments on ENCODE were downloaded and filtered for 8 fold-enrichment and 10^−5^ p-value over the input. Peak files were also downloaded from POSTAR^23^ and used with default filters. Similar to miRNAs, for each RBP we compared the number of actual targets in a regulation group to the number of targets found in a random gene set of the same size to obtain an empirical p-value and z-score (n=100,000 permutations). Benjamini-Hochberg corrected p-values were used for significance.

## DCM disease patient network analysis

Spearman ranked correlation was calculated between the RBP’s log_10_(TPM_RPF_) and target’s log_10_(TE) across patient population (n=30, using all Ribo-seq and RNA-seq matched samples) with the *cor.test* R function. A permutation (n=10,000) test was carried out to determine expected number of correlated pairs (RBP:target) that would be found in a random set. Empirical p-value was calculated and Benjamini-Hochberg correction was applied for multiple testing. RBPs correlating with more pairs than random at FDR > 5% were selected as network hubs. The network was visualized using Cytoscape^60^. To maximise the number of comparisons for the correlation between RBPs and disease severity, we used all 97 patient RNA-seq data. Spearman ranked correlation was calculated between the RBP’s log_10_(TPM_mRNA_) and fibrosis marker’s log_10_(TPM_mRNA_) with the *cor.test* R function. Patient clustering was carried out using hclust R package in default settings based on marker gene expression levels across all 97 patients. Treecut R package was used to obtain four (k=4) levels of severity in fibrosis. Student’s t-test was used to determine significance of the difference between RBP expression for low and high fibrosis severity patients.

## Acknowledgements

The authors would like to acknowledge the technical expertise and support of N.S-J.Ko, B.L.George, M. Wang, J. Schulz and V. Schneider-Lunitz and NGS Team at NHCS. The research was supported by the National Medical Research Council (NMRC) Singapore STaR awards to S.A.C. (NMRC/STaR/0029/2017), the NMRC Central Grant to the NHCS, Goh Foundation, Tanoto Foundation and a grant from the Fondation Leducq. S.S. is supported by the Goh Foundation and the Charles Toh Cardiovascular Fellowship. A.A.W. is supported by the NMRC YIRG (NMRC/OFYIRG/0053/2017). O.J.L.R is supported by NMRC YIRG (NMRC/OFYIRG/0022/2016).

## Author contributions

S.S., S.A.C, N.H. and O.J.L.R conceived and designed the study. E.A., S.V., A.A.W., J.T., C.J.P., L.E.F., J.D., S.B. and G.P. performed in vitro cell culture, cell biology and molecular biology experiments. S.C., S.S., S.R.L., A.A.W., E.A., G.D., and E.G.C. analyzed the data with support from S.v.H. F.W. and P.J.R.B., S.S., S.C., E.A., S.A.C. and O.J.L.R. prepared the manuscript with input from co-authors.

## Competing interests

S.A.C. and S.S. are co-inventors of the patent applications (WO2017103108, WO2017103108, WO 2018/109174, WO 2018/109170). S.A.C. and S.S. are co-founders and shareholders of Enleofen Bio PTE LTD, a company (which S.A.C. is a director of) that develops anti-fibrotics. O.J.L.R is a co–inventor of the patent (WO/2017/106932) and is a co-founder, shareholder and director of Cell Mogrify Ltd, a cell therapy company. All other authors declare no competing interest.

## Data and software availability

The data is available on the gene expression omnibus (GEO id. pending) In order to investigate transcription and translation levels of individual genes we provide a web-resource: http://ribo.systems-genetics.net/

## Supplementary Figures

**Supplementary Figure 1:**
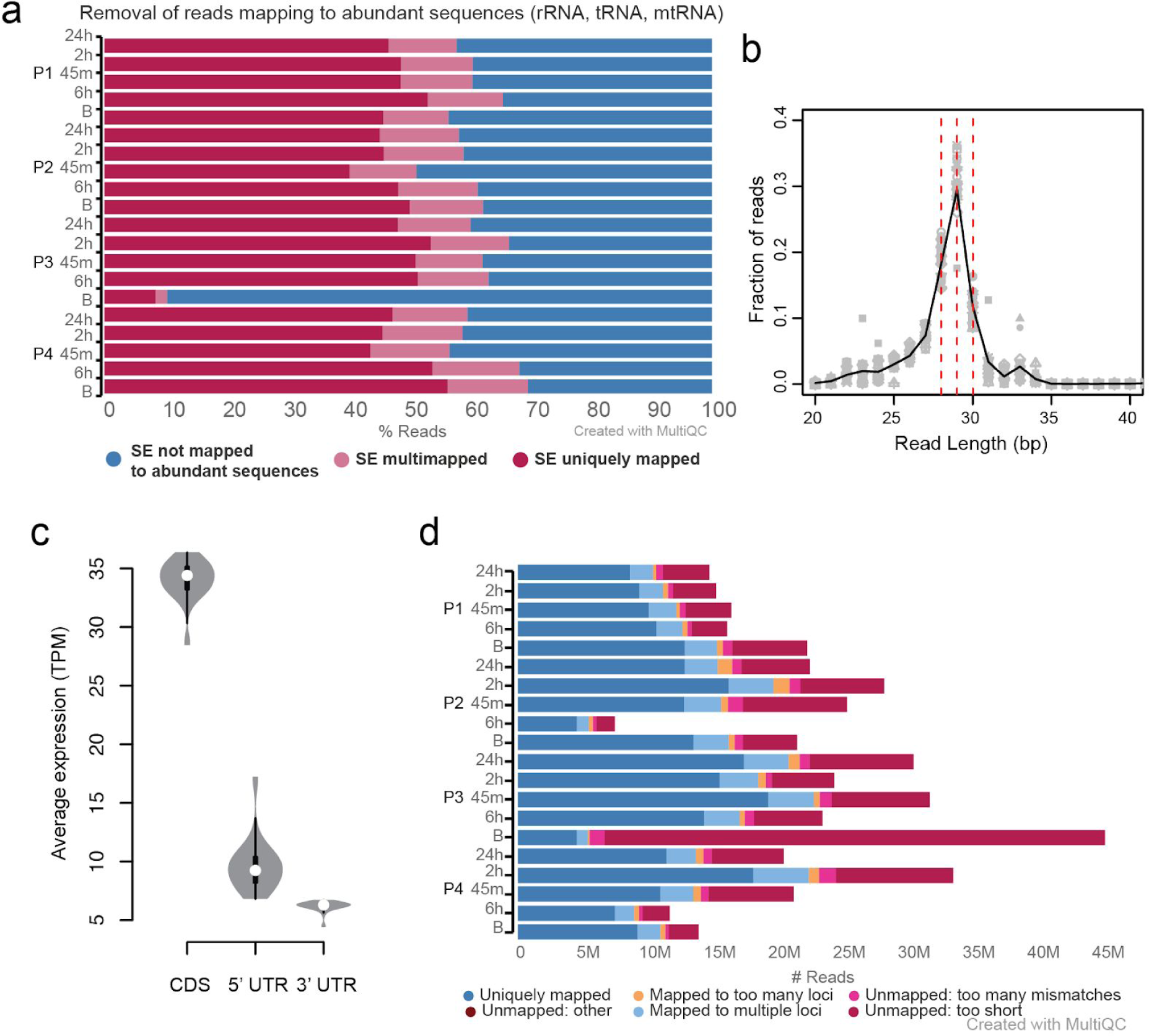
**a.** Percentage of trimmed reads mapping to abundant sequences (ribosomal RNA, transfer RNA, mitochondrial rRNA). **b.** Length distribution of reads after removal of abundant sequences showing 28,29,30 as the predominant lengths of ribosome protected fragments (RPFs). **c.** Average expression (Transcripts per million mapped reads) in each region of protein coding genes showing higher coverage on coding sequence (CDS) compared to the untranslated regions as expected for RPFs. **d.** Number of reads mapping to the human genome (hg38) per sample using STAR aligner.

**Supplementary Figure 2:**
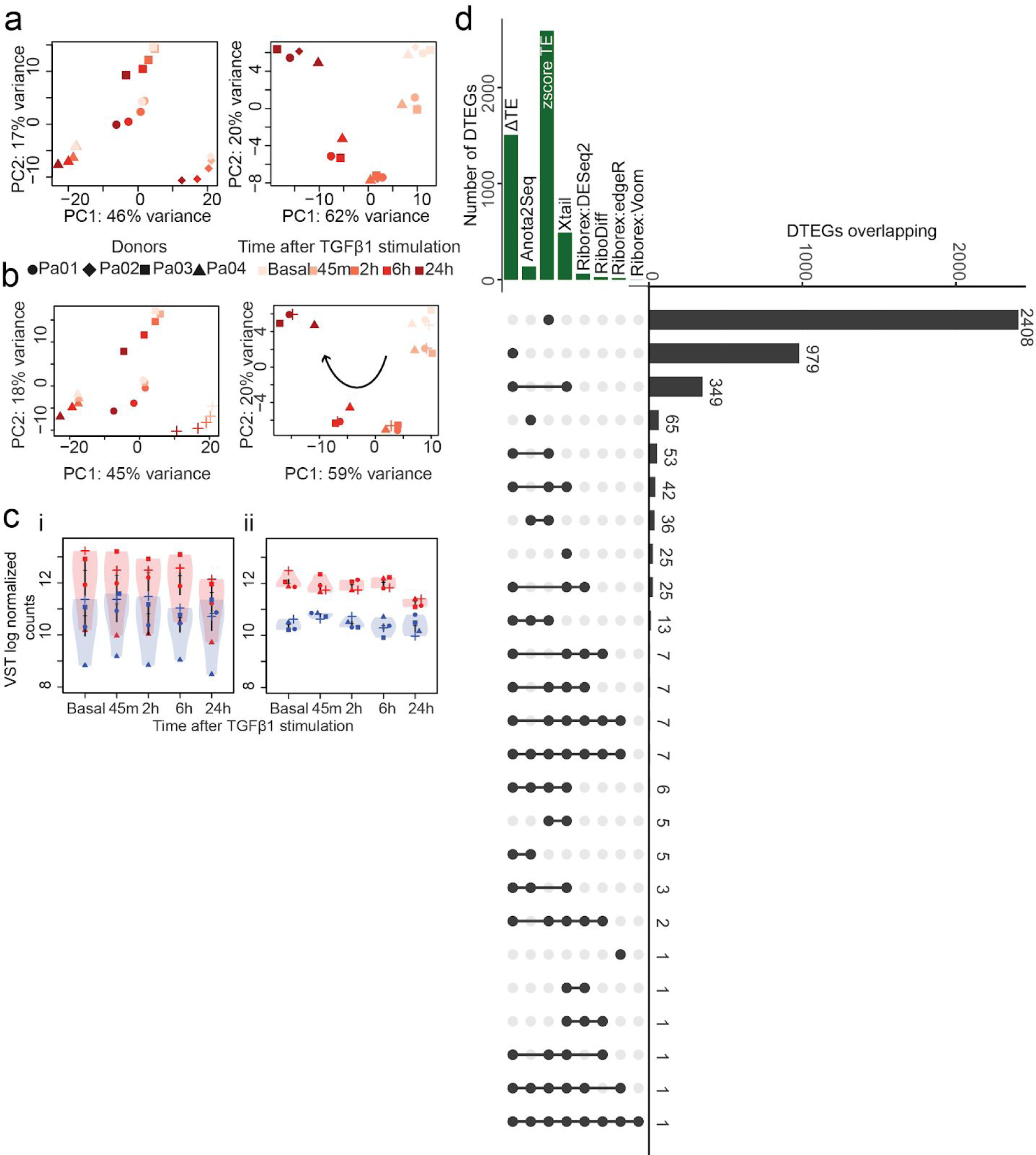
**a.** Principal component analysis (top 500 genes, ranked by variance across samples) of the RNA-seq and **b.** Ribo-seq data across the fibrotic response. PC1 accounts for 46%/45% of the variance in the gene expression and samples grouped by patient. After batch correction, PC1 accounts for 62%/59% of the variance and samples grouped by time-point. **c.** Ribo-seq and RNA-seq counts from Integrin 7A exhibit a strong patient effect, not allowing detection as a DTEG (i), which is ameliorated after batch correction on normalized VST counts (ii). **d.** Number of DTEGs detected and overlap of DTEGs across different published methods in TGFβ1 stimulated primary human fibroblasts.

**Supplementary Figure 3:**
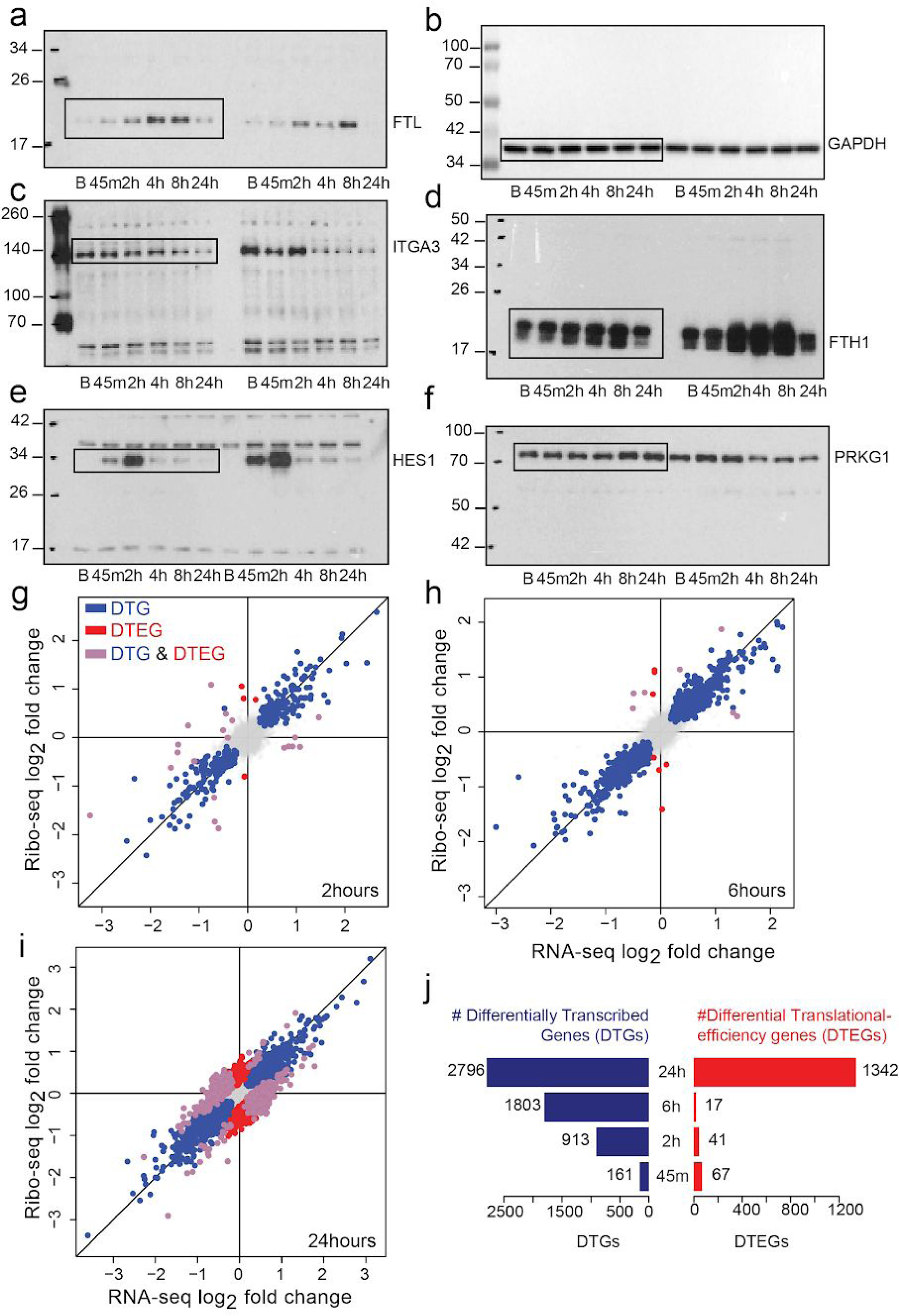
**a-f:** Western blotting of exclusive (FTL (**a**), FTH1 (**c**), ITGA (**d**)), intensified (HES1 (**e**)) and buffered (PRKG1 (**f**)) genes with control (GAPDH (**b**)) in two patients at baseline and 4 time points after TGFβ1 stimulation in primary human cardiac fibroblasts. **g-i:** Log-fold changes in the mRNA and ribosome occupancy at 2 hours (**g**), 6 hours (**h**) and 24 hours (**i**) after TGFβ1 stimulation. DTG: Differentially transcribed genes, DTEG: Differential translational-efficiency genes. **j:** DTGs and DTEGs detected at each time point after TGFβ1 stimulation.

**Supplementary Figure 4:**
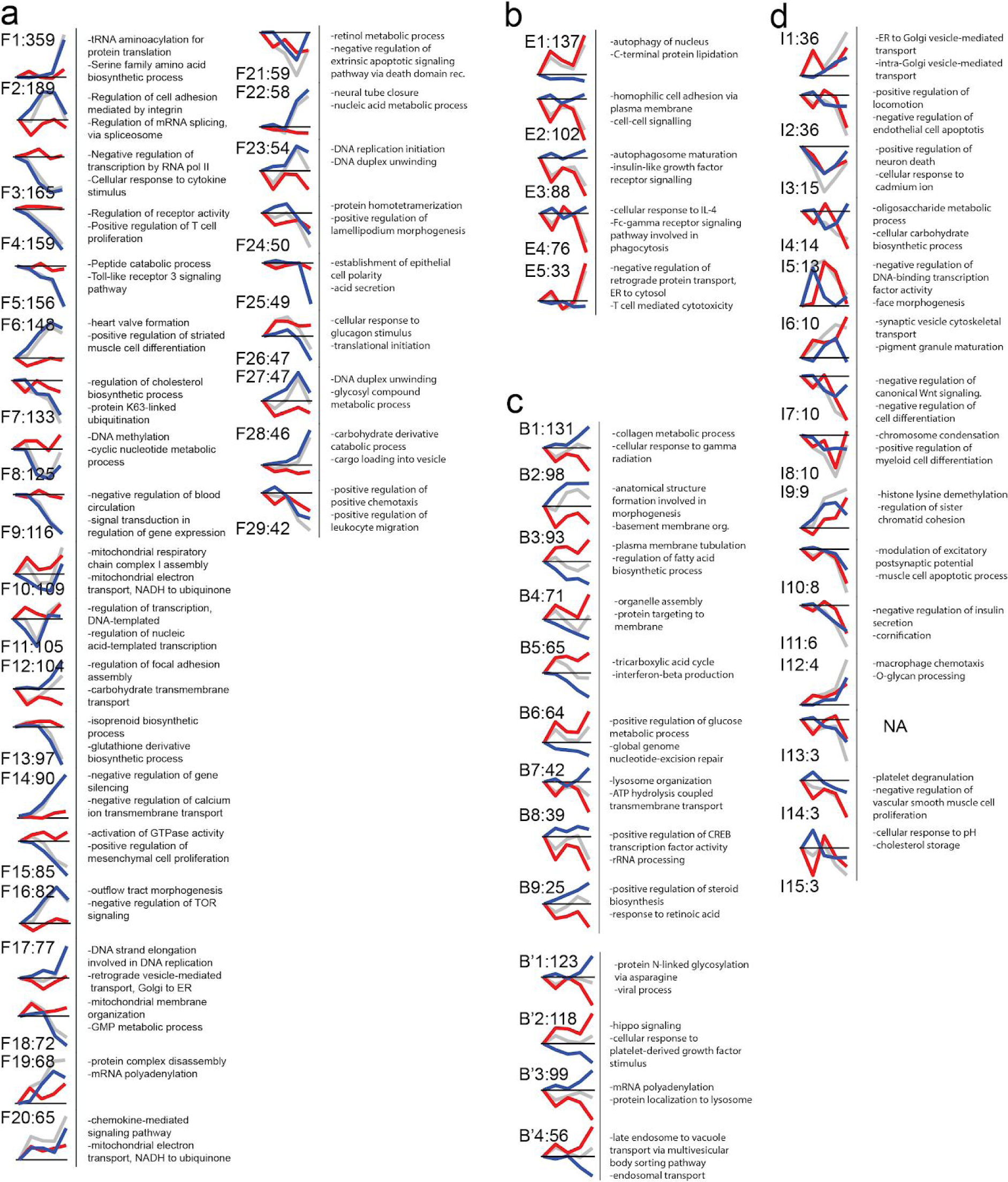
Hierarchical clustering of different regulatory groups including **a.** F: forwarded **b.** E: exclusive **c.** B: buffered, B’: completely buffered (special case) and **d.** I: intensified at the translational level. The cluster IDs are represented as (gene regulatory class)(#):(Number of genes in that cluster). For instance, F1:359 is the first forwarded cluster with 359 genes. The clusters are ordered based on cluster size in each regulatory class.

**Supplementary Figure 5:**
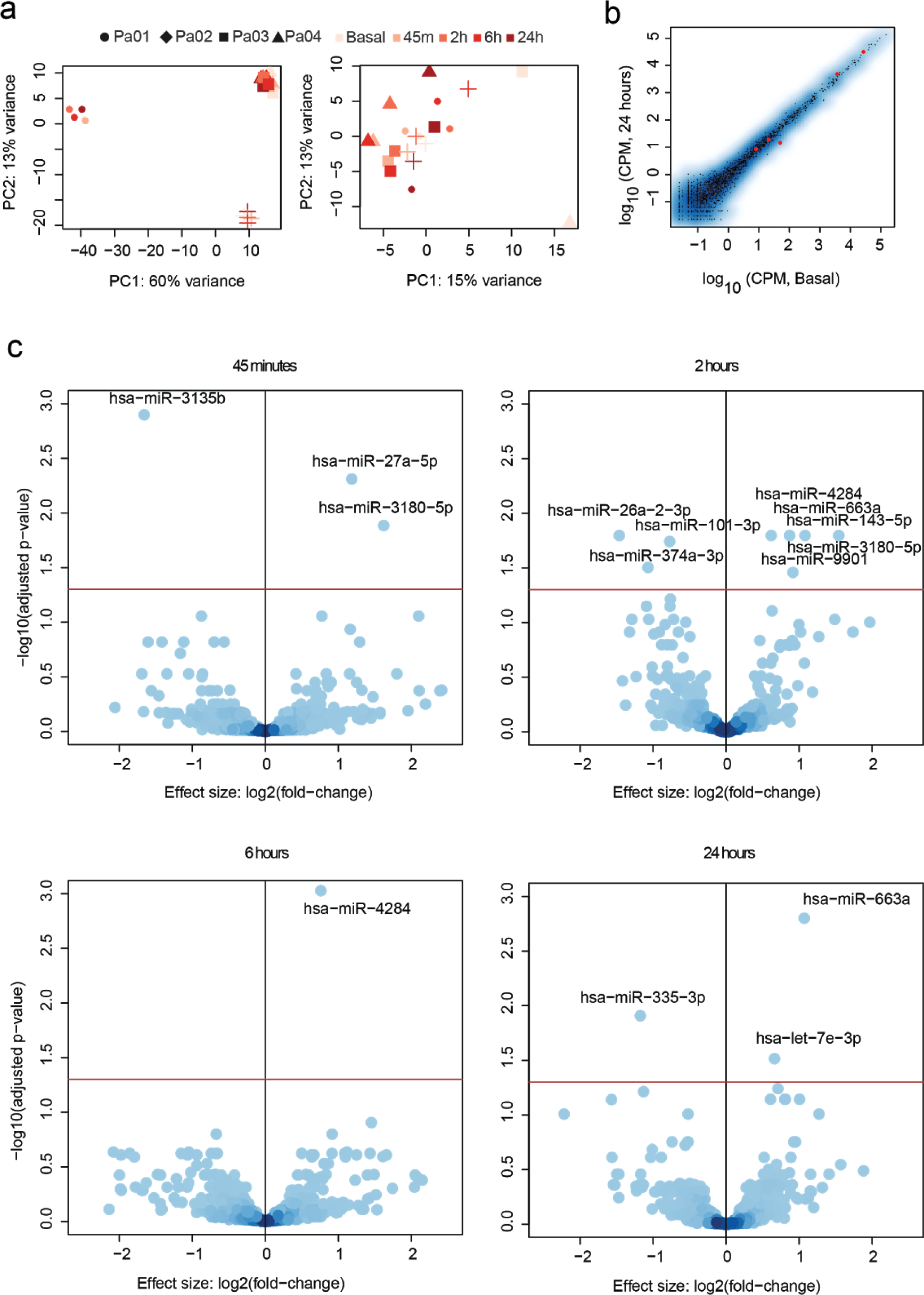
miRNA sequencing of TGβ1 stimulated patient-derived fibroblasts. **a.**Principal component analysis before and after patient effect removal **b.** Counts per million (CPM) for all miRNAs for basal vs 24hrs after TGFB1 stimulation. **c.** Volcano plots for each time point after stimulation, X axis: Log fold change of miRNAs and Y axis: BH corrected p-value. Significantly changing miRNAs are labelled.

**Supplementary Figure 6:**
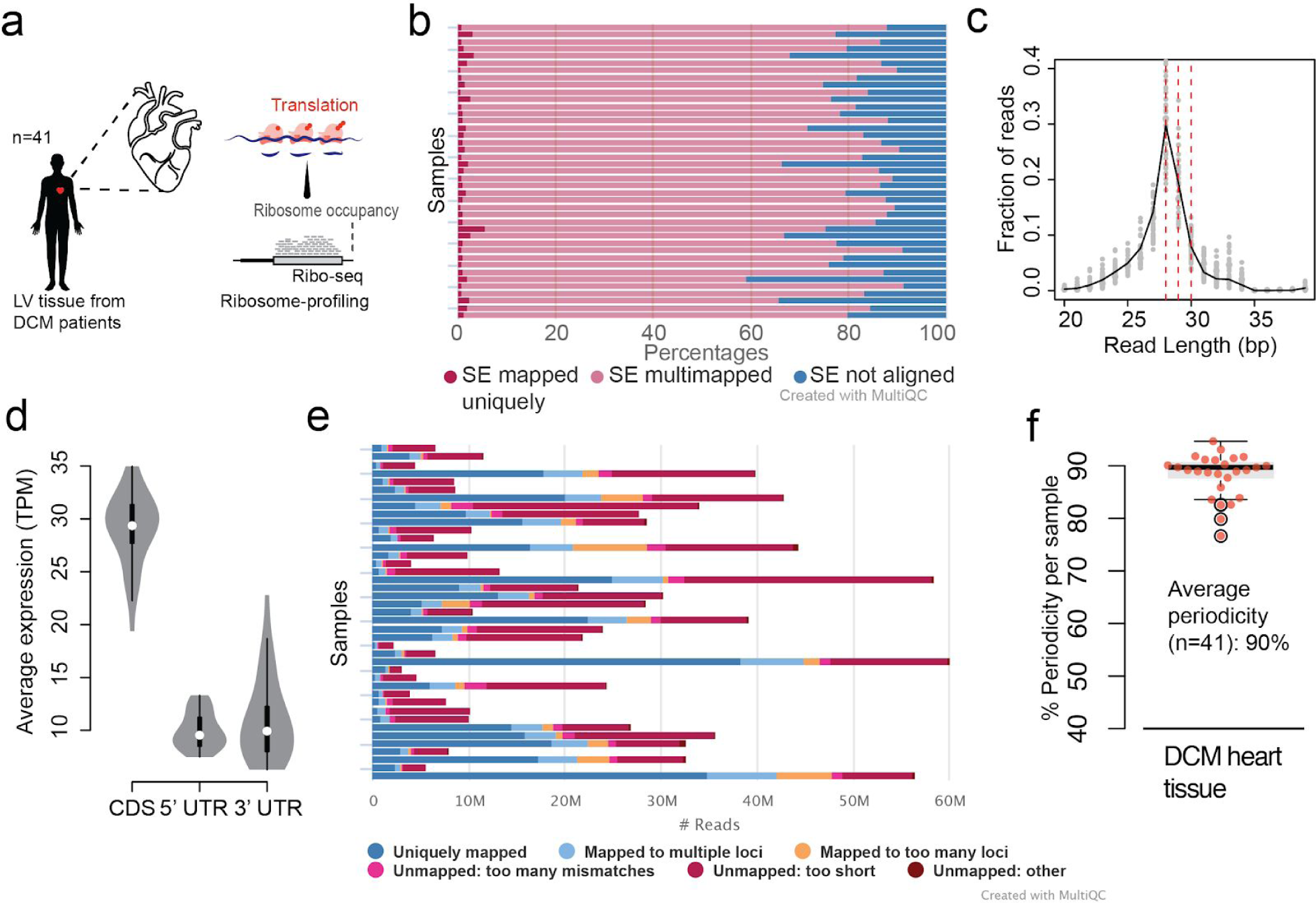
Genome-wide translational profiling of dilated cardiomyopathy patients. **a.** Ribosome profiling of left ventricle tissues from 30 DCM patients **b.** Percentage of trimmed reads mapping to abundant sequences (rRNA, tRNA, mitochondrial rRNA). **c.** Length distribution of reads after removal of abundant sequence mapped reads showing 28,29,30 as the predominant lengths of ribosome protected fragments (RPFs). **d.** Average expression (TPM) in each region of protein coding genes showing higher coverage of coding sequence (CDS) as expected for RPFs. **e.** Number of reads mapping to the human genome (hg38) per sample. **f.** Average periodicity of 90% across all 30 samples.

**Supplementary Figure 7:**
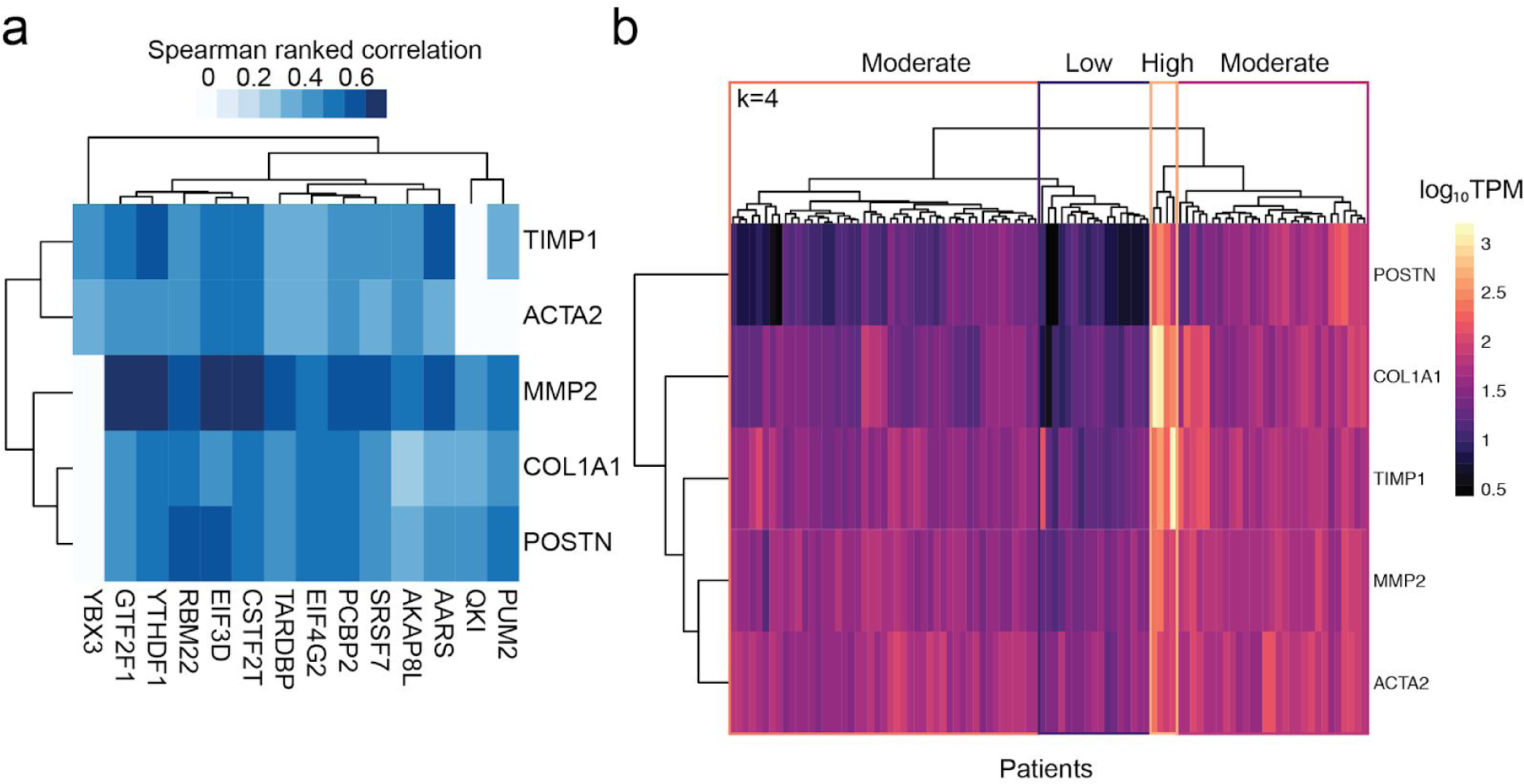
**a.** Gene expression correlation (spearman ranked) of RBPs with fibrosis markers across 97 DCM patients. Only significant correlations based on p-value < 0.05 are shown. **b.** Hierarchical clustering to stratify patients based on fibrosis marker gene expression. Tree was cut into four clusters, resulting in four groups of fibrogenic marker expression severity.

**Supplementary table 1:** Patient information for four male chinese individuals undergoing coronary artery bypass grafting

| Patient | Age |
| --- | --- |
| P1 | 59 |
| P2 | 52 |
| P3 | 57 |
| P4 | 51 |

**Supplementary table 2:** Differentially expressed miRNAs miRNA

| miRNA | Time point<br>vs Basal | Log<sub>2</sub>(Fold change) | p-adjusted |
| --- | --- | --- | --- |
| hsa-miR-27a-5p | 45mins | 1.184228 | 0.004888401 |
| hsa-miR-3135b | 45mins | -1.659315 | 0.001264819 |
| hsa-miR-3180-5p | 45mins | 1.621175 | 0.013024706 |
| hsa-miR-101-3p | 2hrs | -0.772709147 | 0.0181312164 |
| hsa-miR-143-5p | 2hrs | 1.081196984 | 0.01598394 |
| hsa-miR-26a-2-3p | 2hrs | -1.4631523784 | 0.01598394 |
| hsa-miR-3180-5p | 2hrs | 1.5437827151 | 0.01598394 |
| hsa-miR-374a-3p | 2hrs | -1.0687095537 | 0.0313592745 |
| hsa-miR-4284 | 2hrs | 0.6185961048 | 0.01598394 |
| hsa-miR-663a | 2hrs | 0.8701085878 | 0.01598394 |
| hsa-miR-9901 | 2hrs | 0.9156435417 | 0.0348222108 |
| hsa-miR-4284 | 6hrs | 0.759730895 | 0.0009399791 |
| hsa-let-7e-3p | 24hrs | 0.6627012246 | 0.0305164045 |
| hsa-miR-335-3p | 24hrs | -1.1751396741 | 0.012351793 |
| hsa-miR-663a | 24hrs | 1.0699233457 | 0.0015832685 |

**Supplementary table 3:**
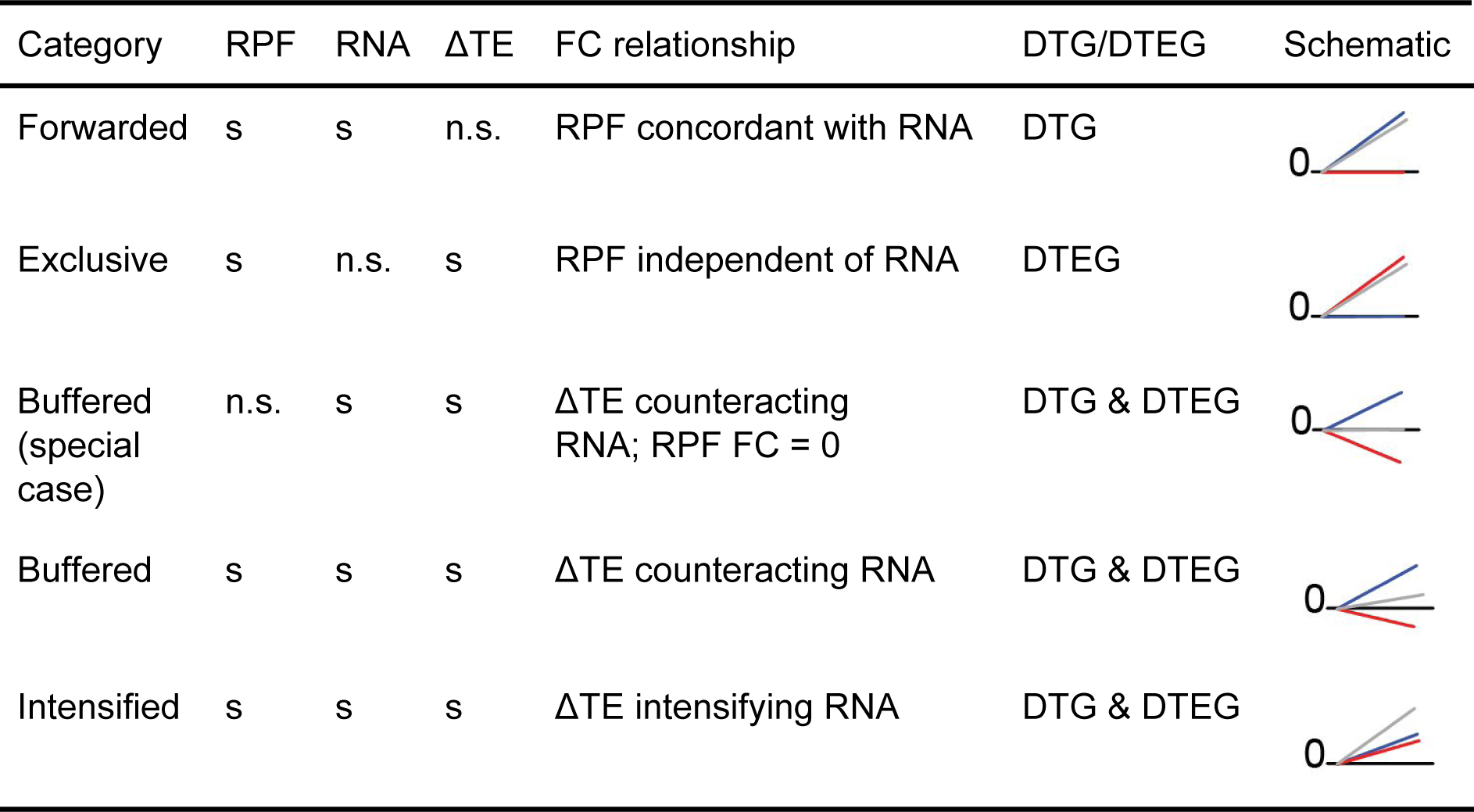
Classification of genes into regulation categories based on significance and fold change relationship. s: Significant changes calculated using DESeq2 with FDR<0.05; n.s: Not significant. FC: Fold change. Concordant: Significant fold changes are in the same direction. Counteracting: Significant fold changes are in the opposite direction. Schematic: RPF: gray, RNA: Blue, ΔTE: Red.

| Category | RPF | RNA | ΔTE | FC relationship | DTG/DTEG | Schematic |
| --- | --- | --- | --- | --- | --- | --- |
| Forwarded | s | s | n.s. | RPF concordant with RNA | DTG |  |
| Exclusive | s | n.s. | s | RPF independent of RNA | DTEG |  |
| Buffered (special case) | n.s. | s | s | ΔTE counteracting RNA; RPF FC = 0 | DTG & DTEG | |
| Buffered | s | s | s | ΔTE counteracting RNA | DTG & DTEG | |
| Intensified | s | s | s | ΔTE intensifying RNA | DTG & DTEG | |

**Supplementary table 4:**
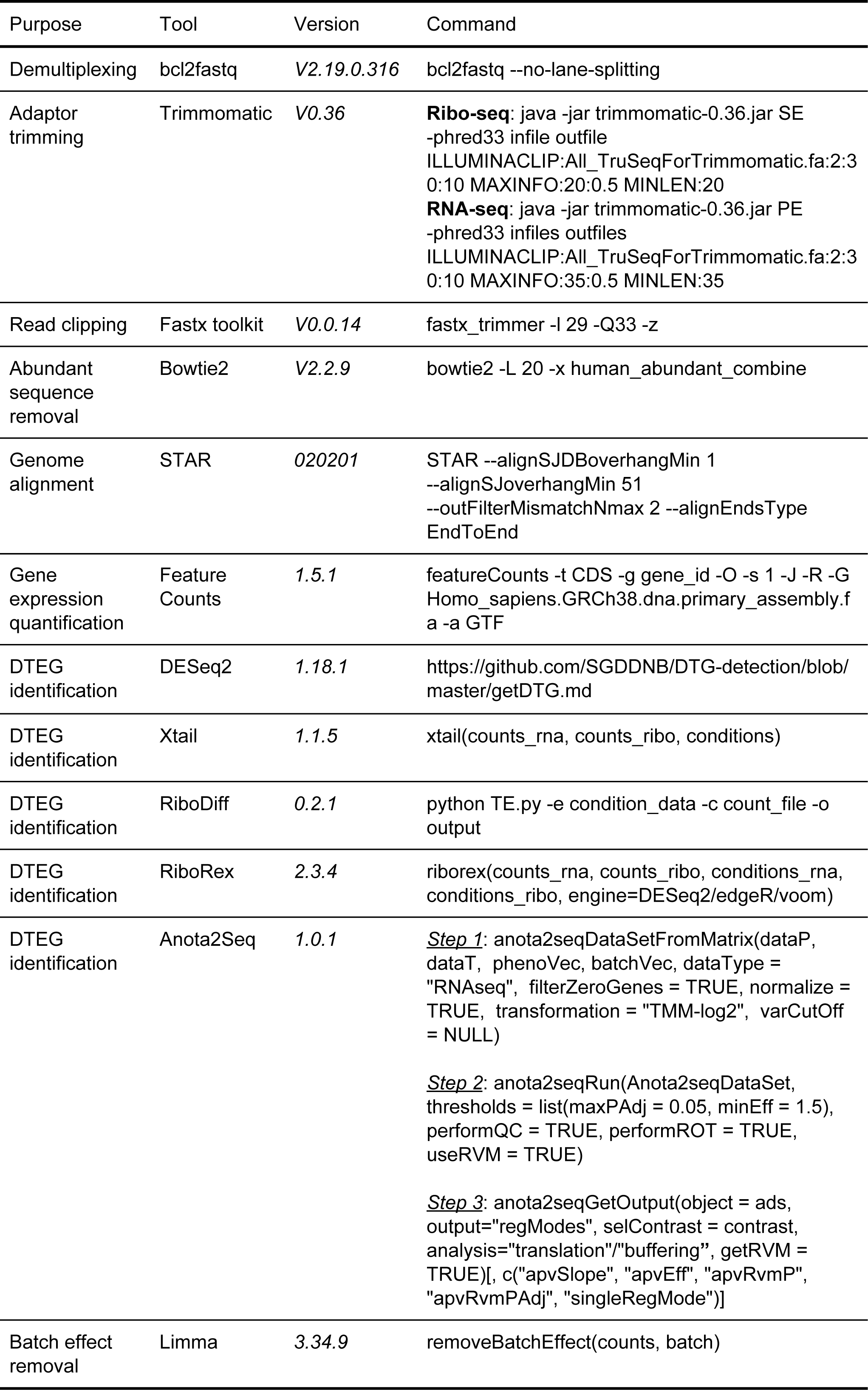
Software and tools used for data analysis

